# A Cross-Trophic Amoebozoan Predator Consumes Trophozoites and Cysts of *Naegleria* and *Acanthamoeba*

**DOI:** 10.64898/2026.05.06.723337

**Authors:** Yonas I. Tekle, Leila N. Plunkett, Angelicia A. Greer, Michael McGinnis

**Affiliations:** Spelman College, 350 Spelman Lane Southwest, Atlanta, Georgia 30314, U.S.A.

**Keywords:** Amoebozoa, *Mayorella*, cross-trophic predation, trophic plasticity, free-living amoebae, cyst predation, microbial food webs, *Naegleria*, *Acanthamoeba*

## Abstract

Protistan predators are key regulators of microbial food webs, yet most are considered to occupy relatively narrow trophic niches. Here, we demonstrate that *Mayorella* spp. (Amoebozoa), isolated from marine and freshwater environments, exhibits exceptional trophic breadth spanning multiple trophic levels. Live-cell imaging revealed predation on bacteria, algae, dinoflagellates, diatoms, flagellates, ciliates, and multicellular prey including rotifers. Large or filamentous prey were engulfed whole or mechanically fragmented during ingestion. Notably, *Mayorella* consumed both trophozoites and cysts of free-living amoebae (*Naegleria* and *Acanthamoeba*), with clear digestion of cyst contents. Dense cultures showed aggregation around large prey and facultative cannibalism. Ingestion of microplastic-like particles occurred without evidence of digestion. Predator cell size and population density increased markedly when feeding on protist or mixed prey relative to bacterial diets, indicating pronounced trophic plasticity. These findings establish *Mayorella* as a broad-spectrum, cross-trophic predator with the capacity to exert top-down effects across microbial food webs and suggest a previously underappreciated role in the suppression of pathogenic free-living amoebae.

## Introduction

Protistan predators exert strong top-down control on microbial communities and play central roles in nutrient regeneration and carbon transfer in aquatic systems (Azam et al. 1983, Anderson 2018, Sherr and Sherr 2002). Microzooplankton can consume a substantial fraction of primary production, typically ∼60–75% of daily phytoplankton output, with values occasionally approaching 90% or higher in highly recycled systems, thereby strongly influencing community composition and biogeochemical fluxes (Calbet and Landry 2004, Landry and Calbet 2004). These interactions extend across diverse environments, from nutrient-poor lakes to deep-sea hydrothermal systems, where protist grazing shapes both bacterial diversity and carbon cycling (Hu et al. 2021). Despite this ecological prominence, protistan predators are often conceptualized as trophically constrained, operating within relatively narrow prey spectra such as bacterivory, algivory, or protistivory (Sherr and Sherr 2002, Pernthaler 2005, Jürgens and Matz 2002), with comparatively few examples of broad cross-trophic predation.

Free-living amoebae (FLA) occupy a critical and complex ecological niche, acting as major consumers of bacteria and small eukaryotes while also serving as reservoirs and evolutionary training grounds for bacterial pathogens (Greub and Raoult 2004, Molmeret et al. 2005, Sherr and Sherr 2002). Genera such as *Acanthamoeba* and *Naegleria* are globally distributed, environmentally abundant, and medically significant, capable of causing serious and often fatal opportunistic infections in humans (Marciano-Cabral and Cabral 2003, Visvesvara et al. 2007). A defining feature of these organisms is their ability to form highly resistant cysts that persist under adverse conditions and contribute to long-term environmental survival (Visvesvara et al. 2007). While amoebae are well recognized as predators of bacteria and hosts of pathogens, their role as predators of other amoebae, particularly targeting resistant cyst stages, remains largely unexplored.

The genus *Mayorella* (Amoebozoa) is widely distributed across marine, freshwater, and terrestrial environments, yet its ecological role remains poorly resolved (Schaeffer 1926, Page 1983a, Page 1983b, Bovee 1961, Cann 1981). Molecular surveys have detected *Mayorella* across diverse habitats, including marine and deep-sea environments, suggesting broad ecological tolerance (Glotova et al. 2018). Classical culture-based studies indicate that some *Mayorella* species grow poorly on bacteria alone and may require eukaryotic prey (Glotova et al. 2018, Page 1983a), hinting at trophic versatility. However, no study has systematically characterized its prey spectrum or ecological function, leaving its position within microbial food webs unclear.

Here, we present evidence from cultures derived from marine and freshwater environments demonstrating that *Mayorella* spp. exhibits an unusually broad prey spectrum spanning multiple trophic levels. Using live-cell imaging, we document predation on bacteria, diverse protists, and small metazoans, as well as the consumption and digestion of both trophozoites and cysts of free-living amoebae, including *Naegleria* and *Acanthamoeba*. We further describe previously unreported behaviors such as mechanical fragmentation of large prey, aggregation during feeding, and cannibalism. These findings suggest that *Mayorella* functions as a cross-trophic predator, linking microbial and microfaunal components of food webs. Notably, its ability to consume both active and cyst stages of opportunistic pathogenic amoebae raises the possibility that *Mayorella* may contribute to the natural suppression of these organisms in environmental systems. Together, these results highlight the potential for certain understudied amoebae to occupy apex-like roles within microbial ecosystems and to influence both community structure and pathogen dynamics.

## Materials and Methods

### Sampling and Culture Establishment

Water and sediment samples used in this study were collected from marine and freshwater environments. Marine samples were obtained from the Whitney Marine Research Center, St. Augustine, Florida, USA (2022), and freshwater samples were collected from Cape Coast, Ghana (2024).

Cultures were established following standard laboratory enrichment protocols for marine and freshwater isolates (Tekle et al. 2022, Tekle et al. 2025). Freshwater samples were maintained in Deer Park water, while marine samples were maintained in 3% artificial seawater. To promote microbial growth and sustain complex microbial communities, sterile grains of rice were added to all cultures.

Mixed microbial communities from nature, including bacteria, algae, protists, and microfauna, were maintained under ambient laboratory conditions for extended periods (2-4 years) without targeted subculturing or selective enrichment. *Mayorella* species were consistently dominant in these natural enrichment cultures. Initial observations of growth and feeding behavior were conducted directly on these unmanipulated communities.

Attempts were made to isolate monoclonal *Mayorella* cultures representing different morphotypes. Most isolation attempts resulted in low growth or culture collapse. However, one freshwater isolate from Ghana was successfully maintained with minimal contamination. This culture persisted in association with a co-occurring *Naegleria* population, which became a primary prey source.

Additional experimental co-cultures were established by introducing *Acanthamoeba castellanii* (CCAP 1501/10) into *Mayorella* cultures. Observations of feeding behavior were conducted across (i) natural mixed communities, (ii) *Mayorella–*bacteria (iii) *Mayorella–Naegleria* co-cultures, and (iv) *Mayorella–Acanthamoeba* co-cultures, as well as behavior of high-density *Mayorella* populations in absence of other preys.

### Microscopy and Imaging

Live observations were conducted using a Nikon inverted phase-contrast microscope. Feeding interactions were monitored over extended periods (minutes to hours) to capture dynamic behaviors. Time-resolved imaging, including still images and live video recordings, was performed using NIS-Elements software, supplemented by smartphone-based imaging. Representative sequences documenting feeding events, including initial contact, engulfment, and digestion, were selected for analysis. Images and videos were processed and assembled into composite figures using Adobe Photoshop and Premiere Pro to visualize feeding behaviors across diverse prey types.

### Prey Identification and Classification

Prey organisms were identified primarily based on morphological characteristics observed during live microscopy. Identified prey included bacteria, algae, diatoms, flagellates, ciliates, amoeboid protists (including *Naegleria*), cyst forms of free-living amoebae, and multicellular microfauna such as rotifers. To confirm the identity of *Mayorella* and associated prey organisms, metabarcoding data from environmental samples were complemented with targeted molecular analyses. DNA was extracted from cultures, and the 18S rRNA gene was amplified using metabarcoding eukaryotic primers. Sequence analyses confirmed affiliation of the amoebae with *Mayorella* spp. of potential new species. Identities of co-cultured protists, including *Naegleria*, were also verified. These sequence data are reported in detail in Tekle et al. (2025, 2026). *Acanthamoeba castellanii* (CCAP 1501/10) used in co-culture experiments was obtained from the Culture Collection of Algae and Protozoa (CCAP).

### Cell size and population density measurements

*Mayorella* cultures were maintained under three feeding conditions: (i) bacterial-only, (ii) free-living amoebae (FLA; e.g., *Naegleria* and/or *Acanthamoeba*), and (iii) mixed-prey communities derived from natural or enrichment cultures. All treatments were incubated under identical environmental conditions and sampled at comparable growth stages to ensure consistency across treatments.

Cell size was quantified from live, actively motile cells using a calibrated Nikon and associated imaging software. For each condition, individual cells were randomly selected (n = 50 per treatment), and the maximum diameter of locomotive morphotypes was recorded. Cell-size ranges were then determined for each feeding condition.

Population density was estimated by direct microscopic counts from well-mixed cultures and expressed as the average number of cells per counted field. All statistical analyses were performed in R version 4.3.1.

## Results

### Broad prey spectrum and handling of structurally complex prey

Observations from both marine and freshwater natural and enrichment cultures revealed that *Mayorella* exhibits an unusually broad prey spectrum spanning multiple trophic levels. In mixed microbial communities and defined microcosms, individual cells actively engulfed bacteria, unicellular and filamentous algae/bacteria, dinoflagellates, diatoms, heterotrophic flagellates, ciliates, and even large parasitic eggs (Fig. 1A–H).

**Figure 1.**
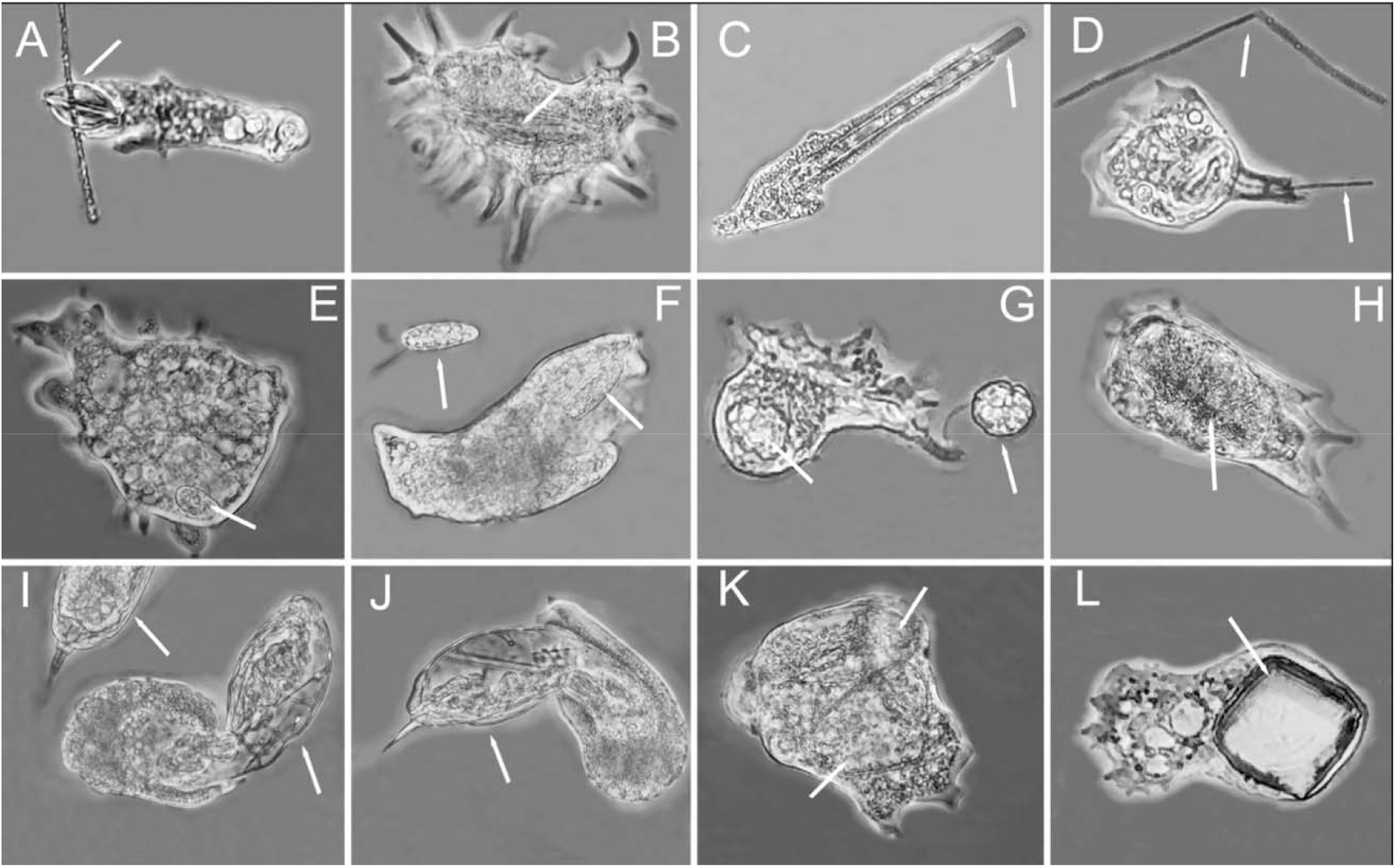
Broad prey spectrum of *Mayorella* across multiple trophic levels. Representative images (A–L) showing predation on diverse prey types: (A–B) ingestion of diatoms; (C) diatom handling and partial digestion; (D) filamentous bacteria; (E) ciliate prey; (F–G) flagellates; (H) parasitic egg; (I–K) rotifer captured and engulfed at successive stages; (L) ingestion of inert particulate material, highlighting the broad trophic range of *Mayorella*. Arrows indicate prey.

Engulfment of smaller unicellular prey typically occurred upon contact, with multiple prey items enclosed simultaneously within distinct food vacuoles. Highly motile protists were captured via cytoplasmic extensions that surrounded the prey (Fig. 1A, F). Once enclosed, prey movement rapidly declined and digestion proceeded within seconds to minutes, with cellular contents progressively assimilated into the cytoplasm (Fig. 2, Supplementary Videos 1–5).

**Figure 2.**
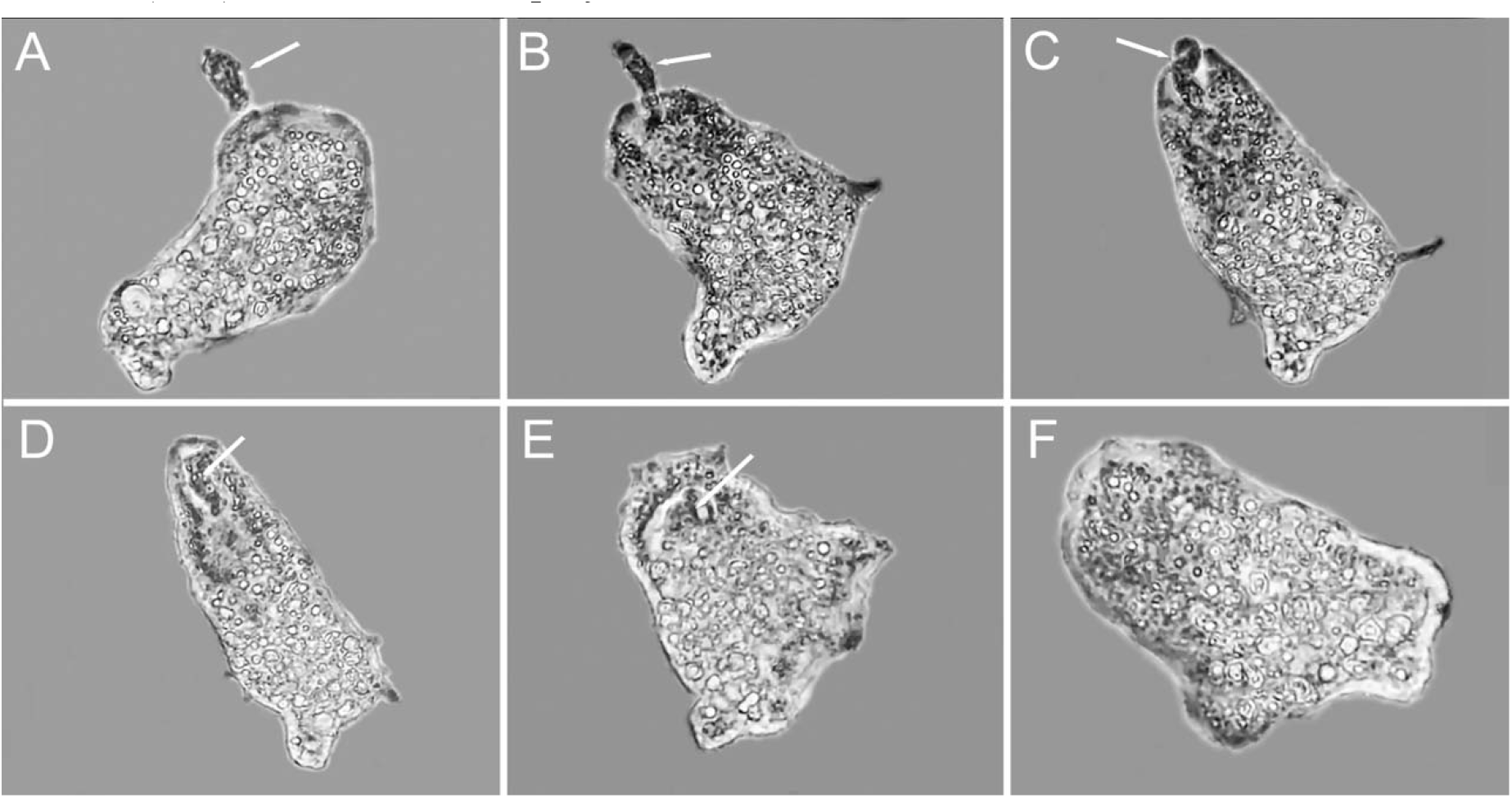
Sequential engulfment of *Naegleria* sp. by *Mayorella*. Representative time series (A–F) showing progressive stages of predation, from initial contact and pseudopodial attachment (A–B), through enclosure and internalization (C–D), to digestion of the prey cell within a food vacuole (E–F). Arrows indicate prey.

Predation extended to multicellular microfauna and structurally protected prey. Rotifers (>400 μm), often two to three times larger than the amoeba, were captured following prolonged pursuit lasting several minutes (Fig. 1I–K; Supplementary Video 1). Successful attacks involved partial enclosure that restricted prey mobility, followed by gradual engulfment. In some instances, multiple amoebae simultaneously attacked (group hunt) a single prey item, although only one individual completed ingestion in moat observed cases (Supplementary Video 1). Feeding primarily targeted internal tissues, leaving the rigid lorica mostly intact.

Prey possessing protective external structures, including diatoms, were similarly susceptible to attack. Amoebae established contact via pseudopodial extensions and progressively enveloped portions or the entirety of the cell. Digestion involved localized breakdown of internal contents, which were extracted into food vacuoles, often leaving behind empty or deformed frustules (Fig. 1C; Supplementary Video 2). Feeding on internally protected prey, such as algae, rotifers, or cysts, involved removal of internal contents; however, the mechanism by which the cell wall or rigid structures were breached was not directly observed. This process may involve perforation or exploitation of existing openings.

Filamentous preys were actively targeted and manipulated during feeding. Filamentous algae, cyanobacterial chains, and filamentous bacteria were mechanically processed prior to or during ingestion. *Mayorella* cells glided along filaments and exerted mechanical force through a barrel-shaped engulfment, forming constrictions that fragmented prey into ingestible segments (Fig. 1D; Supplementary Video 2). These segments were subsequently internalized and digested, demonstrating that *Mayorella* can overcome morphological constraints of prey through active mechanical processing.

### Predation on free-living amoebae and resistant cysts

A key finding was the ability of *Mayorella* to prey on other free-living amoebae (FLA). Trophozoites of both *Naegleria* and *Acanthamoeba* were captured and rapidly engulfed following pseudopodial contact, forming digestive vacuoles in which prey cells were completely degraded within seconds (Figs. 2, 3D; Supplementary Video 3).

**Figure 3.**
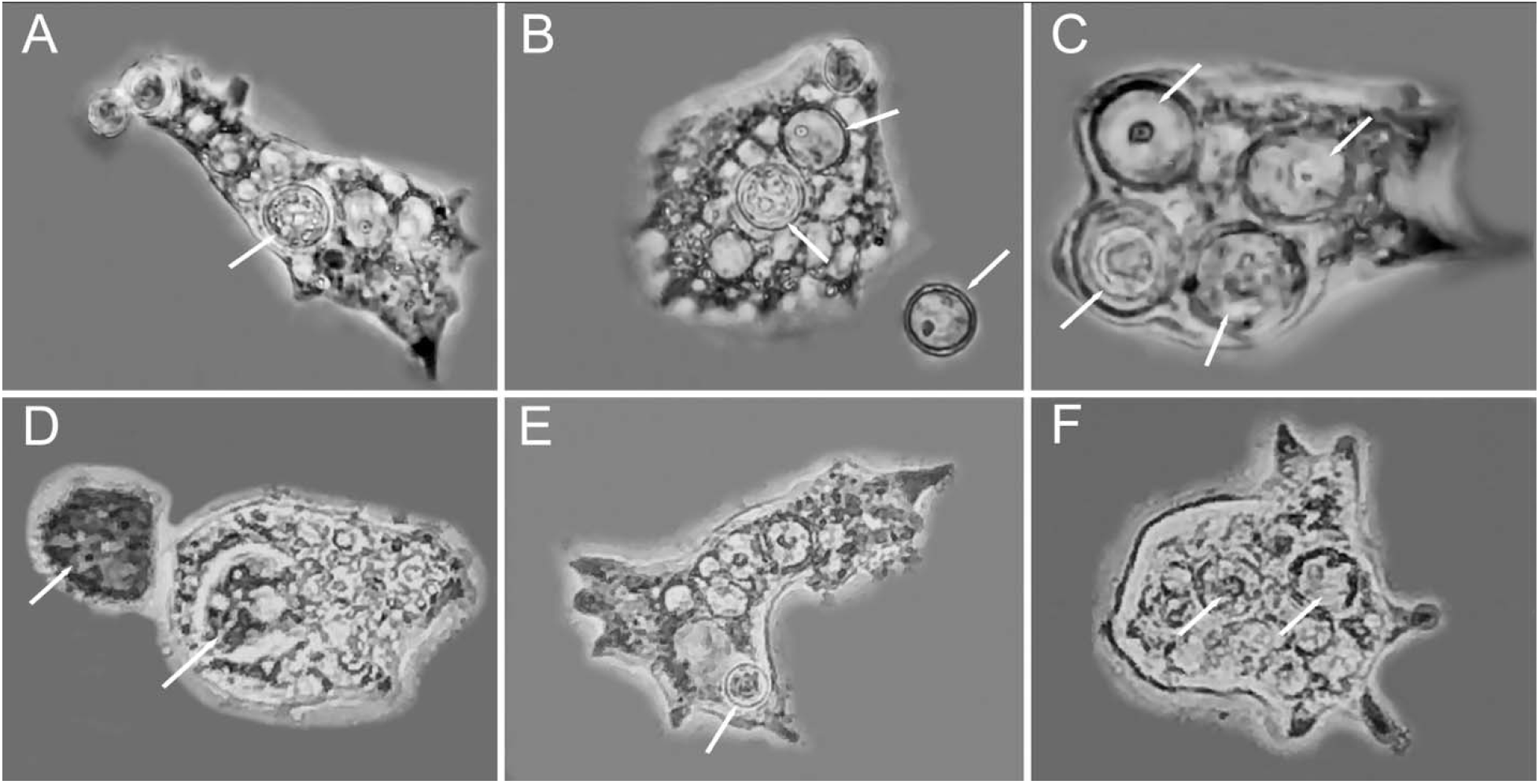
Predation on free-living amoebae and cyst stages by *Mayorella*. (A–C) Sequential engulfment and digestion of *Naegleria* cysts. (D) Ingestion of *Acanthamoeba castellanii* trophozoite. (E–F) Engulfment and digestion of cyst stages. Arrows indicate trophozoite and cysts at different stages of digestion both inside and outside *Mayorella* cells. Digested cysts appear as empty or partially emptied shells with little to no internal content remaining. Arrows indicate prey and FLA cyst at different digestion stages.

At high densities of free-living amoebae (FLA), *Mayorella* spp. were observed grazing on trophozoite stages of FLA (Supplementary Video 3). Under conditions of declining vegetative prey and limited resources, *Mayorella* also consumed cyst stages. Individual cells engulfed one or multiple cysts, which were subsequently observed within food vacuoles undergoing progressive degradation (Fig. 3A–C, F). Time-resolved imaging revealed gradual digestion of cyst contents following engulfment, while cyst walls persisted as empty shells that were later expelled. In many cases, multiple cysts at different stages of digestion were present within a single cell (Fig. 3B, C, F). These observations demonstrate that *Mayorella* can exploit both active and dormant life stages of FLA, including structurally resistant cysts.

### Density-dependent interactions: aggregation and cannibalism

Under conditions of abundant or large prey, *Mayorella* cells occasionally aggregated around a single prey item. Two or more individuals were observed simultaneously feeding on the same microorganism (Supplementary Videos 1, 2, 4), suggesting opportunistic co-feeding or competitive aggregation. In most cases, feeding interactions ultimately resulted in a single individual retaining the prey.

At high cell densities and under prey-limited conditions, cannibalistic behavior was observed (Supplementary Video 4). Larger cells engulfed smaller conspecifics following prolonged pursuit, including recently divided daughter cells. These events were primarily observed in confined, high-density cultures with limited external prey and were uncommon under mixed-community conditions. Together, these observations indicate density-dependent behavioral plasticity, including both aggregation and intraspecific predation.

### Ingestion of inert particulate matter

In addition to biological prey, *Mayorella* cells were observed engulfing non-living particles, including microscopic sand grains and microplastic-like particulates present in cultures (Fig. 1L; Supplementary Video 5). These particles were internalized into food vacuoles but remained structurally intact and were not digested. Ingested particles were retained transiently and later expelled, indicating that *Mayorella* is a size-based, largely non-selective phagocyte at the point of ingestion, with selectivity occurring during intracellular processing.

### Growth and size plasticity across feeding conditions

*Mayorella* exhibited pronounced variation in cell size and population density across feeding conditions (Fig. 4). Under bacterial-only conditions, cells remained relatively small (40–60 μm) and occurred at low densities (∼10 cells per counted field). In the presence of free-living amoebae (FLA), cell size increased (60–120 μm), accompanied by a marked rise in population density (∼90 cells per counted field). The largest cells were observed in mixed-prey communities (100–200 μm), which also supported the highest densities (∼120 cells per counted field).

**Figure 4.**
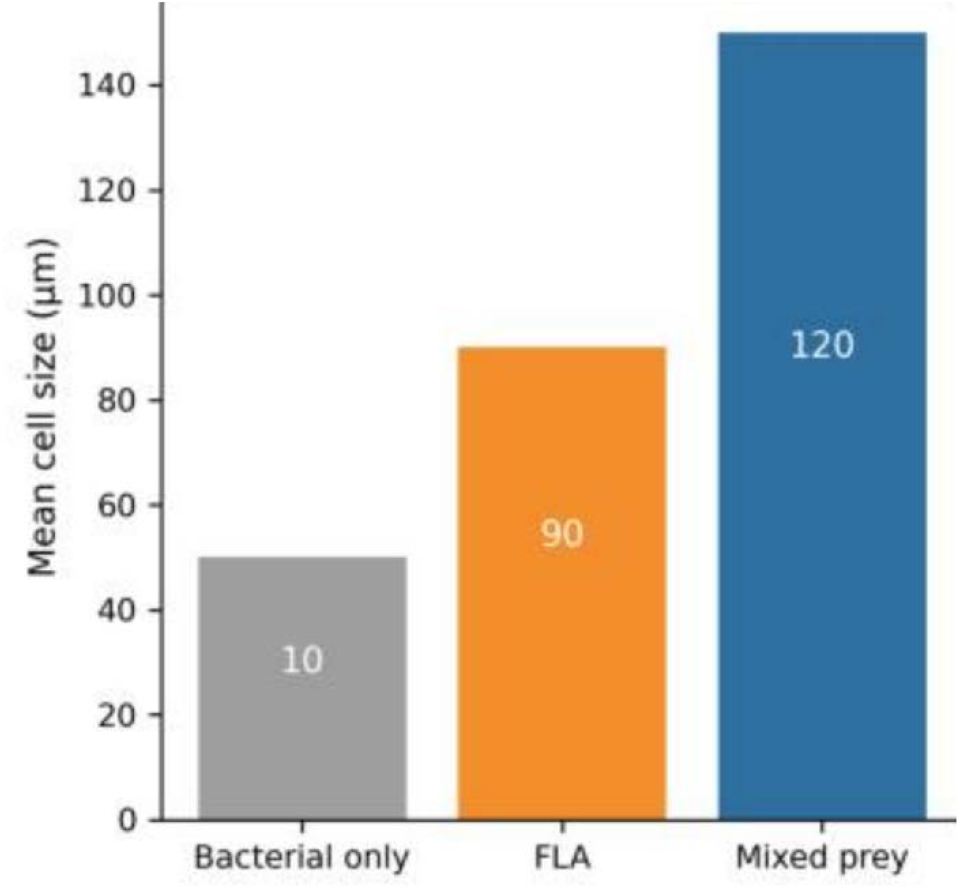
Cell size and population density of *Mayorella* across feeding conditions. Bar heights represent mean cell size (µm) under bacterial-only, free-living amoebae (FLA), and mixed-prey conditions. Numbers within bars indicate average population density (cells per counted field): 10, 90, and 120, respectively. Both cell size and population density increase with prey complexity.

These patterns indicate strong trophic plasticity, with both growth and population expansion closely linked to prey complexity and availability.

## Discussion

Our observations indicate that *Mayorella* functions as a cross-trophic generalist predator, with a prey spectrum spanning multiple levels of the microbial food web. This breadth is notable, as most protistan predators are typically constrained to narrower feeding niches such as bacterivory or algivory (Landry and Calbet 2004, Pernthaler 2005, Sherr and Sherr 2002). By linking bacterial grazing, algal predation, and consumption of other protists and microfauna, *Mayorella* integrates multiple trophic pathways and is well positioned to influence community structure through multi-level predation. Protists play central roles in nutrient regeneration and carbon transfer (Anderson 2018, Azam et al. 1983, Hu et al. 2021), yet relatively few taxa have been shown to operate simultaneously across these functional domains. In this context, *Mayorella* may represent an ecological hub that couples bacterial, protistan, and microfaunal compartments within a single predatory framework.

A notable feature of *Mayorella* is its apparent ubiquity and persistence across environments. It was consistently recovered from both marine and freshwater systems and frequently emerged as a dominant amoeboid form in enrichment cultures (Tekle et al. 2026). Such recurrence suggests that *Mayorella* can persist across diverse conditions and may represent a stable component of microbial communities. Similar patterns of persistence and enrichment dominance have been reported for other free-living amoebae (FLA), which can maintain long-term populations through flexible feeding strategies and environmental resilience (Smirnov et al. 2011, Smirnov and Goodkov 1999). In this context, the consistent presence of *Mayorella* across diverse communities, coupled with its broad trophic capacity, suggests a potentially underrecognized role in structuring microbial population dynamics and may position it as an indicator of active trophic interactions within complex microbial ecosystems.

The ability to consume resistant stages of pathogenic amoebae represents a particularly significant ecological function. Free-living amoebae such as *Naegleria* and *Acanthamoeba* include species responsible for severe and often fatal infections, and their persistence in natural and engineered systems is largely driven by highly resistant cyst stages (Samba-Louaka 2023, Sriram et al. 2008). These cysts can survive disinfection, desiccation, and prolonged environmental exposure, forming long-term reservoirs of infection. Our observations demonstrate that *Mayorella* not only consumes trophozoites but also actively engulfs and degrades cysts. The progressive digestion of cyst contents, coupled with the release of empty cyst walls, indicates a feeding strategy capable of overcoming structural defenses. Similar extraction-based feeding was observed across multiple prey types, including metazoans with rigid coverings and algal cells, suggesting a generalized capacity to exploit protected biological material. Together, these findings point to a previously unrecognized mechanism by which predatory amoebae may influence the persistence and transmission potential of pathogenic FLA in environmental systems.

This capability positions *Mayorella* as a previously unrecognized biological agent that may reduce both active and dormant stages of FLA populations. Given that FLA can harbor intracellular bacterial pathogens (Greub and Raoult 2004, Balczun and Scheid 2017, Tekle et al. 2021), predation by *Mayorella* may have cascading effects on pathogen persistence and transmission. While the ecological and applied implications require further investigation, these findings support a framework in which protistan predators contribute to the natural regulation of environmentally persistent pathogens. Comparisons with other protistan predators further highlight the distinctiveness of *Mayorella*. Many microbial predators exhibit specialization in prey type or ecological niche. For example, *Actinophrys sol* can dominate extreme environments and consume metazoans (Bell et al. 2006), testate amoebae can engage in coordinated predation (Geisen et al. 2015), vampyrellids specialize in algal feeding (Hess 2017), and *Oxyrrhis marina* plays a role in regulating algal populations (Goldman et al. 1989). In contrast, *Mayorella* combines bacterivory, algivory, protistivory, metazoan predation, and cyst consumption within a single organism. This multi-modal trophic strategy suggests a level of ecological flexibility that remains underappreciated among amoebozoans.

Consistent differences in cell size and population growth across feeding conditions further support the ecological significance of this trophic breadth. Access to protist and mixed prey resulted in substantial increases in both cell size and population density relative to bacterial-only conditions (Fig. 4), indicating that prey quality directly influences growth potential. These observations are consistent with broader evidence that prey quality and diversity shape protistan growth and biomass production (Mitra et al. 2016, Anderson 2018), and suggest that trophic flexibility enables *Mayorella* to exploit fluctuating resource landscapes, reinforcing its role as a dynamic regulator within microbial communities.

Together, these findings support a model in which *Mayorella* functions as an apex-like amoeboid predator within microbial ecosystems. Rather than occupying a single trophic niche, it traverses multiple trophic levels, integrates diverse energy pathways, and targets both active and resistant life stages of prey. This perspective expands current understanding of protistan ecological roles and highlights the importance of incorporating trophic generalists into conceptual and quantitative models of microbial food webs (Worden et al. 2015).

Importantly, the breadth of feeding strategies observed here suggests an underlying molecular and cellular versatility that remains to be explored. The ability to process structurally diverse prey, including algal isolates, cysts and multicellular organisms, implies a complex combination of enzymatic and mechanical capabilities. Ongoing and future genomic and transcriptomic analyses will provide critical insights into the molecular basis of this predatory capacity and may reveal novel pathways involved in prey recognition, digestion, and structural degradation.

Taken together, these findings suggest that shifts in microbial community composition, particularly under conditions of increased turbidity, nutrient loading, and reduced dissolved oxygen, **may be associated with the presence and persistence of opportunistic pathogens and ecologically impactful taxa**. This highlights the value of microbial eukaryotic communities as indicators of both ecosystem condition and potential health-related risks in freshwater systems (Sagova-Mareckova et al., 2021).

## 6. Conclusion

This study demonstrates that microbial eukaryotic communities in the Budamu and Suruwi freshwater systems are structured by a combination of trophic interactions, environmental filtering, and physicochemical gradients. Dominance of SAR and Obazoa taxa reflects active microbial food web dynamics, while diversity patterns indicate the influence of both hydrological connectivity and localized environmental conditions. The integration of metabarcoding and culture enrichment approaches revealed complementary aspects of microbial diversity, highlighting the presence of both abundant and low-frequency taxa, including ecologically significant and potentially pathogenic lineages. Physicochemical analyses suggest that variables such as turbidity, dissolved oxygen, and total dissolved solids play important roles in shaping community structure, although no single factor alone explains observed patterns. Instead, microbial assemblages appear to be influenced by multiple interacting environmental drivers. These findings underscore the value of combining molecular and morphological approaches for freshwater microbial ecology and highlight the potential of microbial eukaryotes as indicators of ecosystem condition and environmental change. Importantly, this study contributes to addressing the limited availability of molecular-based assessments of microbial eukaryotic diversity in freshwater systems in West Africa and provides a framework for future ecological monitoring and biodiversity studies in the region.

## Acknowledgments

This work was supported by the National Science Foundation Excellence in Research (EiR) Award #2401946 and the Simons Foundation Fellow Award (SFA-23-5) to Y.I.T. We are also grateful to Saron Ghebezadika and Christon Jairus Marquez Racoma for help with laboratory work.

## SUPPLEMENTARY VIDEO CAPTIONS (1–5) (*YouTube Links to the Videos are provided for review purposes*)

**Supplementary Video 1**. Predation on rotifers and diverse protists by *Mayorella*. Time-resolved video sequences showing pursuit, attack, and engulfment of a rotifer, as well as flagellates and ciliates, by *Mayorella*. The footage includes instances of multiple amoebae interacting with a single rotifer prey item, illustrating coordinated or competitive feeding behavior.

***YouTube Link*** : https://youtu.be/l3wFAmtWMNY

**Supplementary Video 2**. Predation on diatoms and filamentous bacteria. *Mayorella* feeding on diatoms, including extraction of internal contents following partial enclosure, and mechanical severing of filamentous bacteria into ingestible segments.

***YouTube Link*** : https://youtu.be/9eyMpcmoQPI

**Supplementary Video 3**. Predation on free-living amoebae. Engulfment and digestion of trophozoites and cysts of *Naegleria* and *Acanthamoeba castellanii* by *Mayorella*.

***YouTube Link*** : https://youtu.be/NIV0aCh-4Fk

**Supplementary Video 4**. Cannibalism and aggregation behavior. Intraspecific predation showing engulfment of conspecific cells, along with aggregation of multiple *Mayorella* individuals under high-density or prey-limited conditions.

***YouTube Link*** : https://youtu.be/c5Oi_xmGCh8

**Supplementary Video 5**. Ingestion of particulate material. Engulfment and subsequent release of inert particulate matter, demonstrating non-selective ingestion and lack of digestion of non-biological material.

***YouTube Link*** : https://youtu.be/1J-cJPyUXK8

